# Early urinary protein changes in the tumor-forming process of the NuTu-19 lung metastasis

**DOI:** 10.1101/519215

**Authors:** Jing Wei, Na Ni, Wenshu Meng, Yuhang Huan, Youhe Gao

## Abstract

The early detection of cancer is essential for effective intervention. Urine, which lacks homeostatic control, has been used to reflect early systemic changes in subcutaneous cancer models. However, urine has not been used to predict whether tumors will be formed in animal models. In this study, a cancer model was established by the tail-vein injection of 2 million NuTu-19 tumor cells. Approximately half of the rats formed lung metastatic carcinoma tumors. Urine samples were randomly selected from 4 tumor-forming rats and 4 non-tumor-forming rats on day 0/12/27/39/52 and analyzed by label-free proteomic quantitative analysis. Compared to day 0, a total of 180 and 118 urinary proteins in tumor-forming and non-tumor-forming rats, respectively, showed significant changes. Functional enrichment analysis of the differential proteins in tumor-forming rats revealed similar events during cancer metastasis cascades and tumor progression, such as the migration of tumor cell lines, coagulation system, TGF-β signaling, the STAT3 pathway and the adhesion of myeloid cells and alveolar macrophages. The differential proteins in non-tumor-forming rats were associated with agranulocyte/granulocyte adhesion and diapedesis, glutathione biosynthesis, IL-12 signaling and vitamin metabolism. After validation by parallel reaction monitoring (PRM) targeted proteomics quantitative analysis, a total of 22 urinary proteins showed significant changes in the early phase of the lung tumor formation process, 5 of which have been associated with the mechanisms of lung cancer, while 3 have been suggested as lung cancer biomarkers. Another 14 urinary proteins showed significant changes in the early phase in non-tumor-forming rats. Our results indicate that urine proteins could differentiate early tumor-forming and non-tumor-forming status.

## Introduction

Cancer biomarkers are measurable changes associated with the pathophysiological processes of cancers that have the potential to diagnose cancer, monitor cancer progression, predict cancer recurrence, and assess treatment efficacy[1]. Unlike blood, urine is a noninvasive and attractive biomarker resource that has been underestimated. Without homeostatic mechanisms, urine can accumulate all changes throughout the body and has the potential to reflect early and small pathological changes[2, 3]. However, the urine proteome is easily affected by various factors, such as age, gender, diet and medication, especially in clinical patients[3]. To solve this issue, animal models facilitate minimizing these influential factors. Animal models can establish the direct relationship between disease and the corresponding changed urinary proteins[4]. The urine proteome has already been used to reflect early changes in various diseases before clinical manifestation in different animal models. For example, i) urine proteins could enable the early detection of cancer at the early onset of tumor growth before a tumor mass was palpable as well as monitoring its progression[1]; ii) urine proteins could enable the early detection and monitoring of both disease progression and treatment efficacy in the BLM-induced pulmonary fibrosis rat model[5]; iii) a panel of differential urinary proteins observable before clinical symptoms may provide sensitive early biomarkers for the early diagnosis of astrocytoma[6]; iv) early urinary candidate biomarkers were detected before the alanine aminotransferase and aspartate transaminase changes in the serum and before fibrosis was observed upon hematoxylin and eosin (HE) and Masson staining in a rat thioacetamide-induced liver fibrosis model[7]; v) urinary proteins can enable the early detection of Alzheimer’s disease before amyloid-β plaque deposition, which may provide an opportunity for intervention[8]; and vi) urinary proteins can reflect early changes caused by chronic pancreatitis[9]. However, no studies on whether urine can differentiate between the presence and absence of tumor formation have been reported. If we found that some animals formed tumors while others did not when establishing cancer animal models, could the difference be reflected in the urine?

Tail-vein injection is usually used to establish cancer lung carcinoma metastatic animal models[10–12]. In this study, a tail-vein injection rat model was established by the injection of two million NuTu-19 tumor cells. Urine samples were collected on days 0, 12, 27, 39, and 52. The numbers of tumor-forming and non-tumor-forming rats were recorded after two months. This study was designed to identify urine proteome changes in tumor-forming and non-tumor-forming rats. The technical flowchart is presented in Figure 1.

**Figure 1:**
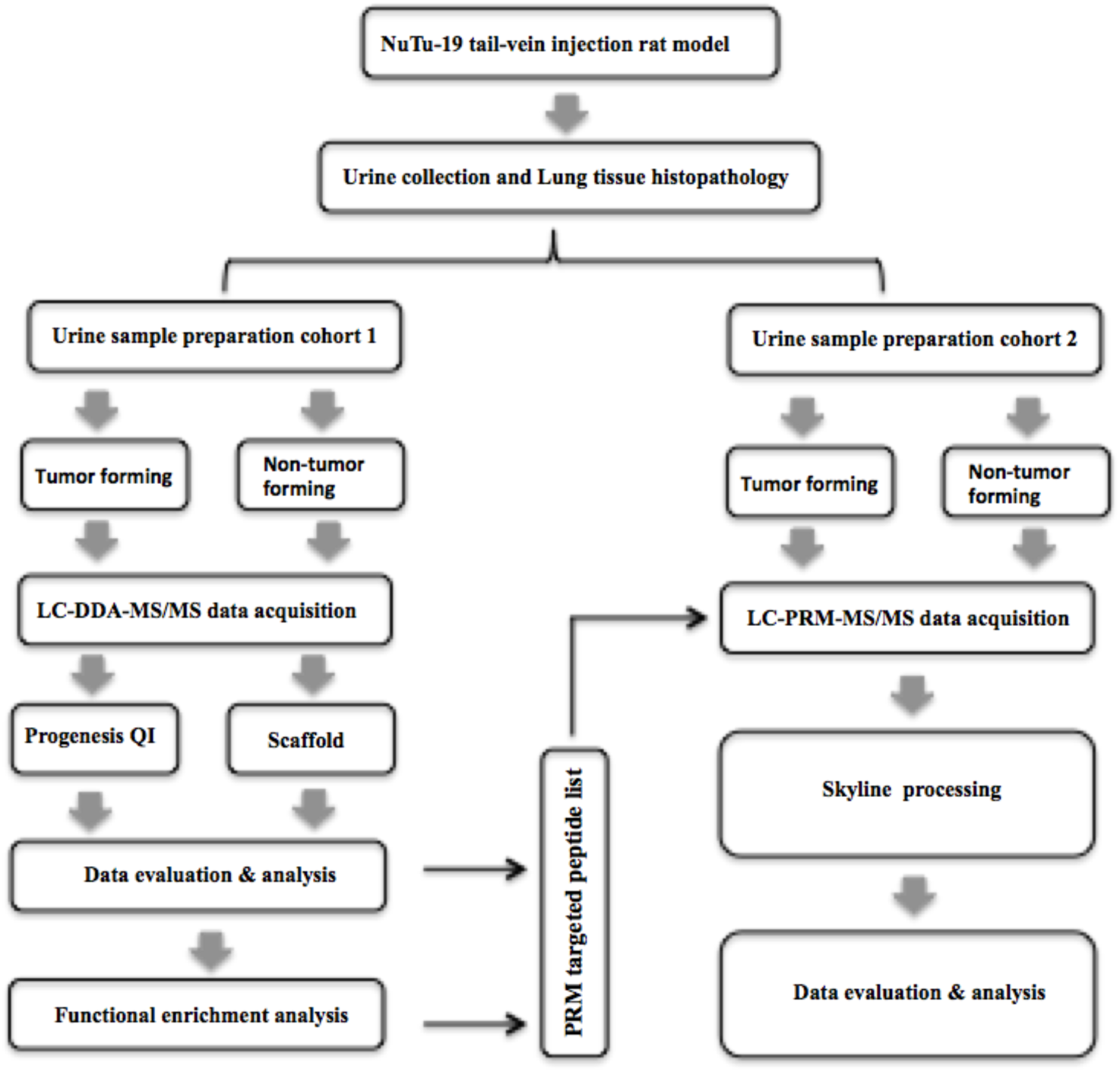
Workflow for the early detection of tumor-forming and non-tumor-forming process. After the tail-vein injection of NuTu-19 cells, urine samples were collected on days 12, 27, 39 and 52. The urine proteome was identified using liquid chromatography coupled with tandem mass spectrometry (LC-MS/MS) analysis. Then, parallel reaction monitoring (PRM)-targeted proteomics quantitative analysis was used to validate candidate biomarkers for the early detection of tumor-forming and non-tumor-forming processes.

## Materials and Methods

### Ethics statement

Male Wistar rats (150 ± 20 g) were purchased from Beijing Vital River Laboratory Animal Technology Co., Ltd. All animals were maintained with a standard laboratory diet under controlled indoor temperature (21±2°C), humidity (65–70%) and 12 h/12 h light–dark cycle conditions. All animal protocols governing the experiments were approved by Peking Union Medical College (Approval ID: ACUC-A02-2014-008). This study was performed according to the guidelines developed by the Institutional Animal Care and Use Committee of Peking Union Medical College. All efforts were made to minimize suffering.

### Experimental Design

The tail-vein NuTu-19 injection rats were established by the tail-vein injection of NuTu-19 ovarian cancer cells, which were purchased from MEIXUAN Biological Science and Technology Ltd. (Shanghai, China). NuTu-19 ovarian cancer cells were cultured at 37°C in RPMI_1640 (Corning) supplemented with 10% fetal bovine serum (GIBCO), 100 IU/mL penicillin (Macgene) and 100 µg/mL streptomycin (Macgene) in a humidified atmosphere of 5% CO_2_ in air (Thermo Fisher Scientific, Inc., Waltham, MA, USA). NuTu-19 tumor cells were stained with 0.4% trypan blue to estimate the cell viability and then counted using a hemocytometer. Their viability was approximately 95% before the tail-vein injection. Cells in the logarithmic growth phase were used for the experiment.

Male Wistar rats were randomly divided into two groups: the tail-vein NuTu-19 injection group (n = 24) and the control group (n = 8). The tail-vein NuTu-19 injection group was injected with 2×10^6^ viable NuTu-19 cells in 100 *µ*L of PBS. The control group was injected with the same volume of PBS via the tail vein. Animals were anesthetized with sodium pentobarbital solution (4 mg/kg) before the tail-vein injection.

The self-controlled experiment was conducted in two phases: for the discovery phase, differential protein identification was performed in four tumor-forming rats and four non-tumor-forming rats; for the validation phase, urine samples were obtained from another four tumor-forming rats and three non-tumor-forming rats. Each rat provided five urine samples.

### Histopathology

For lung histopathology, three rats in the control group and nine rats in the NuTu-19 tail-vein injection group were randomly sacrificed on days 35, 45, and 52 (n=3 each time point) by using an overdose of sodium pentobarbital anesthetic. Another fifteen NuTu-19 tail-vein injection rats were sacrificed on day 60. The whole lung tissue was fixed in 4% formalin fixative and embedded in paraffin. Then, paraffin sections (4 µm thick) were stained with hematoxylin and eosin (HE) to reveal the metastatic tumors.

### Urinary protein extraction

Urine samples were collected from the NuTu-19 tail-vein injection group on days 0, 12, 27, 39, and 52. Each rat was placed in a metabolic cage overnight for 12 h to collect urine without any treatment. After collection, urine samples were stored immediately at −80°C.

Urine samples were centrifuged at 12,000 g for 30 min at 4°C. The supernatants were precipitated with three volumes of ethanol at −20°C overnight. After centrifugation at 12,000 g for 30 min, the pellets were resuspended in lysis buffer (8 mol/L urea, 2 mol/L thiourea, 50 mmol/L Tris, and 25 mmol/L DTT) at 4°C for 2 h. The supernatants were then centrifuged at 4°C and 12,000 g for 30 min. Protein samples were measured by using the Bradford assay and stored at −80°C for later use.

### Tryptic digestion

Proteins were digested with trypsin (Trypsin Gold, Mass Spec Grade, Promega, Fitchburg, Wisconsin, USA) by using filter-aided sample preparation methods[13]. Briefly, 100 *µ*g of proteins was loaded onto 10-kD cutoff filter devices (Pall, Port Washington, NY) and washed twice with UA (8 M urea in 0.1 M Tris-HCl, pH 8.5) and 25 mmol/L NH_4_HCO_3_ at 14,000 g for 40 min at 18°C. Then, each urinary protein was denatured with 20 mM DTT at 37°C for 1 h and alkylated with 50 mM iodoacetamide (IAA) at room temperature for 40 min in the dark. After being washed twice with UA and 3 times with 25 mmol/L NH_4_HCO_3_, the denatured proteins were resuspended in 25 mmol/L NH_4_HCO_3_ and digested with trypsin (enzyme to protein ratio of 1:50) at 37°C for 14 h. The collected peptide was desalted using Oasis HLB cartridges (Waters, Milford, MA) and then dried by vacuum evaporation (Thermo Fisher Scientific, Bremen, Germany).

### Data acquisition by MS

Two types of data acquisition, data-dependent acquisition (DDA) and the parallel reaction monitoring (PRM) targeted proteomics quantitative method, were performed on an Orbitrap Fusion Lumos Tribrid mass spectrometer (Thermo Fisher Scientific, Waltham, MA) coupled with a Thermo EASY-nLC 1200 HPLC system. Digested peptides were redissolved in 0.1% formic acid to a concentration of 0.5 *µ*g/*µ*L. The digested samples were loaded into a trap column (75 µm × 2 cm, 3 µm, C18, 100 Å) at a flow rate of 0.25 µl/min and separated with a reversed-phase analytical column (75 µm × 250 mm, 2 µm, C18, 100 Å). Peptides were eluted with a gradient extending from 5%–30% buffer B (0.1% formic acid in 80% acetonitrile) for 60 min /120 min with the DDA method and for 120 min with the PRM method.

Liquid chromatography (LC)-DDA-MS/MS was performed on 20 samples of 4 tumor-forming rats and 4 non-tumor-forming rats. One microgram of each peptide from an individual sample was loaded on the trap column. The MS data were acquired using the following parameters: i) survey MS scans were acquired in the Orbitrap using a range of 350–1550 m/z with the resolution set to 120,000; ii) MS/MS scans per full scan with top-speed mode (3 s) were selected for collision-induced dissociation fragmentation with the resolution set to 30,000 in Orbitrap; iii) dynamic exclusion was employed with a 30 s window to prevent the repetitive selection of the same peptide; iv) the normalized collision energy for HCD-MS2 experiments was set to 30%; and v) the charge-state screening was set to +2 to +7 and a maximum injection time was 45 ms. Each sample was analyzed with two technical replicates.

Twenty urine samples from another four NuTu-19 tumor-forming rats and fifteen urine samples from three non-tumor-forming rats were chosen for PRM analysis. The initial targeted LC-PRM-MS/MS was performed by DDA to define peptide retention times. Prior to individual sample analysis, peptides were pooled (2 µg of each sample) for LC-MS/MS analysis to build a spectrum library with 6 runs. A total of 900 ng of pooled or individual peptide was separated on a reversed-phase analytical column. MS data were acquired using the following parameters: full scans (m/z 350–1550) were acquired with a resolution of 60,000; PRM scans (m/z 200– 2000) were run at a resolution of 30,000; the retention time window was set to ±2 min; targeted peptides were isolated using a 1.6 m/z window; 30% HCD of normalized collision energy was used; and the maximum injection time was 60 ms.

### Database searching and label-free quantitation

Label-free raw MS/MS data files were searched using Mascot software (version 2.5.1, Matrix Science, London, UK) against the SwissProt rat database (released in February 2017, containing 7992 sequences). The search parameters were set as follows: the parent ion tolerance was 10 ppm, and the fragment ion mass tolerance was set to 0.02 Da. The carbamidomethylation of cysteine was set as a fixed modification, and the oxidation of methionine was considered a variable modification. The specificity of trypsin digestion was set for cleavage after K or R, and two missed trypsin cleavage sites were allowed.

The raw MS data files of NuTu-19 tumor-forming rats were processed using Progenesis software (version 4.1, Nonlinear, Newcastle upon Tyne, UK) for label-free quantification[14]. Features with only one charge or with more than five charges were excluded from the analyses. Protein abundance was calculated from the sum of all unique peptide ion abundances for a specific protein in each run. The normalization of abundances was required to allow comparisons across different sample runs by this software. For further quantitation, all peptides (with Mascot score >30 and P<0.01) of an identified protein were included. Proteins identified by at least one peptide were retained. Only high-confidence peptide identifications with a false discovery rate (FDR) ≤ 0.01 were imported into Progenesis LC-MS software for further analysis.

The raw MS data files of NuTu-19 non-tumor-forming rats were processed using Scaffold software (version 4.7.5, Proteome Software Inc., Portland, OR). Peptide identifications were accepted at an FDR of less than 1.0%, and protein identifications were accepted at an FDR less than 1.0% with at least two unique peptides. Comparisons across different samples were performed after normalization of the total spectra. Spectral counting was used to compare protein abundances at different time points according to a previously described procedure[15, 16].

### Parallel reaction monitoring (PRM) data analysis

In the discovery phase, candidate protein biomarkers were identified using the label-free quantitative proteomics method. Common differential proteins identified in tumor-forming and non-tumor-forming rats were excluded from the subsequent PRM analysis. Differential proteins identified on days 12 and 27 in tumor-forming and non-tumor-forming rats were validated by PRM.

The raw pooled MS/MS data files were searched using Thermo Proteome Discover 2.1.0.81 against the SwissProt rat database (released in February 2017, containing 7992 sequences) with precursor and fragment mass tolerances of 10 ppm and 0.02 Da, respectively. Other parameters were set as follows: trypsin digested; maximum missed cleavage sites of 2; oxidation (+15.995 Da) of methionine as a dynamic modification; and carbamidomethylation (+57.021 Da) of cysteine as a static modification. The protein FDR, which was determined by a target-decoy search strategy, was set to 1%. The Skyline software (*Version 3.6.1 10279*) was used to build a spectrum library and filter peptides for PRM analysis[17]. Two to six peptides of each targeted protein were selected using the following criteria: (i) identified in the untargeted analysis with Mascot score >30 and P<0.01, (ii) digested by trypsin [KR/P] with a maximum of 2 missed cleavages, (iii) 8-18 amino acid residues, (iv) exclusion of the first 25 N-terminal amino acids, and (v) carbamidomethyl (C) and oxidation (M) as the structural modifications. Only unique peptides of each protein were used for the subsequent targeted quantitation. The retention time (RT) segment was set to 4 min for each targeted peptide with its expected RT in the center based on the pooled sample analysis. For the tumor-forming groups, 91 differentially expressed proteins identified in the discovery phase were selected for PRM validation. After further optimization, 78 proteins with 237 peptides were ultimately used for validation by PRM-targeted proteomics (Supplementary Table S1). The technical reproducibility of the PRM assay was assessed; among the targeted peptides, 219 peptides had CV values less than 20% (Supplementary Figure S1). For the non-tumor-forming group, 44 differentially expressed proteins identified in the discovery phase were used for the subsequent PRM analysis. Finally, 42 proteins with 205 peptides could be targeted (Supplementary Table S2). When the technical reproducibility of the PRM assay was assessed, among these 205 targeted peptides, 201 peptides had CV values less than 20% (Supplementary Figure S2).

Individual peptide samples (900 ng of each sample) were then analyzed by PRM assays. The transition settings in Skyline were as follows: precursor charges were set to +2, +3, and +4, ion types to y,p,b, the product ions from ion 3 to last ion, the ion match tolerance to 0.02 m/z, the number of product ions was set to 6, and the min dotp was set to 0.7. Each protein was quantitated using the summation of the fragment area from its corresponding transitions. Prior to the statistical analysis, the summation of fragment area was performed by log2 transformation. The differential proteins were identified using one-way ANOVA, and significance was set at a *P*-value < 0.05.

### Gene Ontology and Ingenuity Pathway Analysis

All proteins found to be differentially expressed in tumor-forming and non-tumor-forming rats were assigned gene symbols using DAVID[18] and analyzed by Gene Ontology (GO) based on the biological process, cellular component and molecular function categories. The biological pathway analysis and disease/function analysis of differential proteins analyzed at four time points were performed by IPA software (Ingenuity Systems, Mountain View, CA, USA)

### Statistical analysis

Average normalized abundance or spectral counts of each sample were used for statistical analysis. The levels of proteins identified in the tumor-forming or non-tumor-forming group on days 12, 27, 39 and 52 were compared with their levels on day 0. Differential proteins were selected with the following criteria: fold change ≥1.5 or ≤ 0.67; confidence score ≥ 200; *P* < 0.05 by two-sided, unpaired t-test; protein spectral counts or normalized abundance from every rat in the high-abundance group greater than those in the low-abundance group; and the average spectral count in the high-abundance group ≥ 4. The P-values of group differences were adjusted by the Benjamini & Hochberg method[19]. Group differences resulting in adjusted *P-values* < 0.05 were considered statistically significant. All results are expressed as the mean ± standard deviation.

## Results and Discussion

### Characterization of NuTu-19 tumor<u>-</u>forming and non-tumor<u>-</u>forming rats

Thirty-two male Wistar rats (150 ± 20 g) were divided into the control group (n=8) and the NuTu-19 tail-vein injection group (n=24). The rats in the control group had normal daily activities with shiny hair, while some of the NuTu-19 tail-vein injection rats showed piloerection in the late stages. Fifteen rats were killed on day 60. Eight rats showed obvious lung tumors, while the other seven rats did not. The lung histopathology is presented in Figure 3. The average body weight of the tumor-forming rats was significantly lower than that of the control rats until day 47 (p<0.05), while the average body weight of the non-tumor-forming group was lower than that of the control group but without significance (Figure 2).

**Figure 2:**
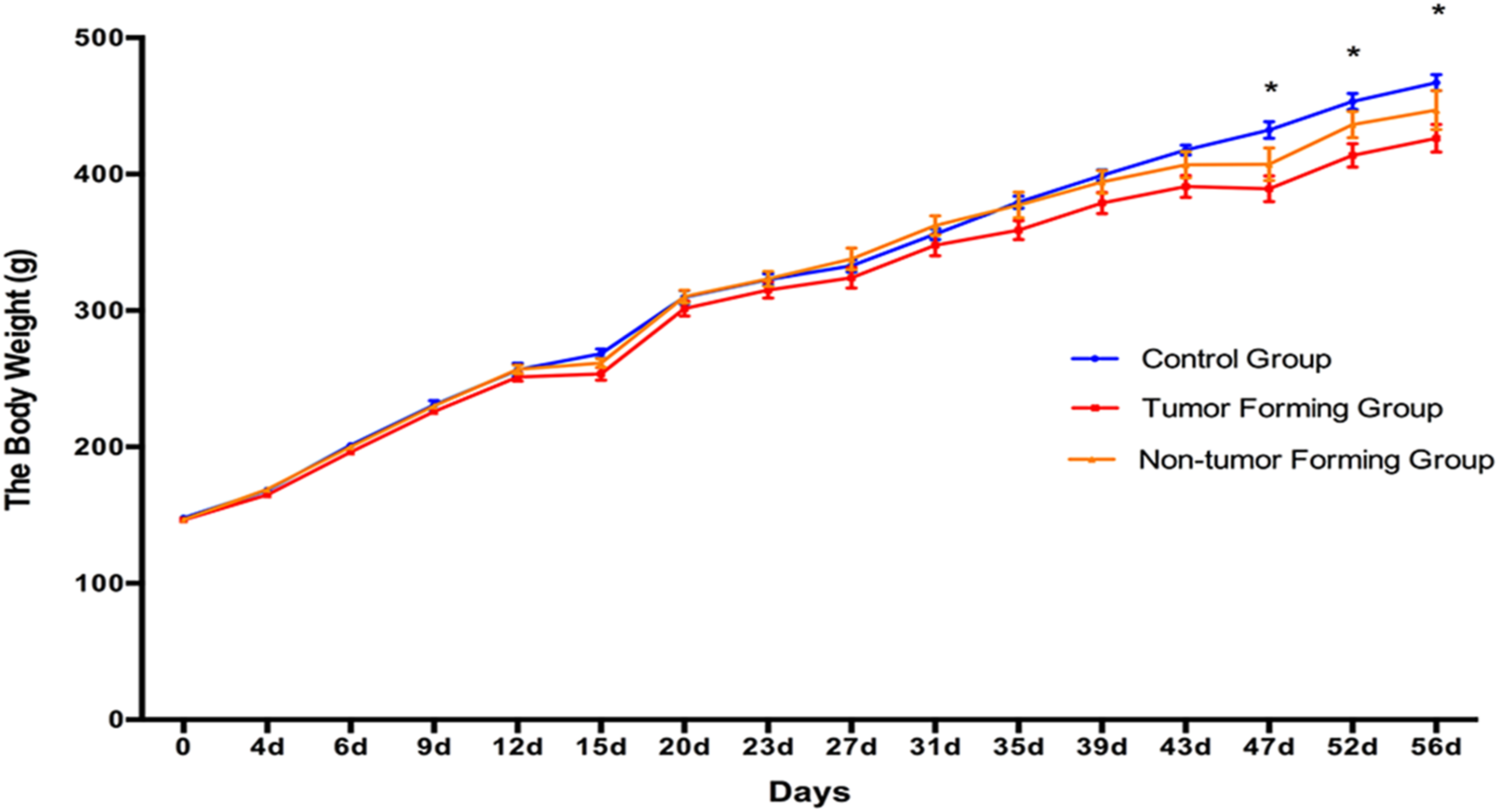
The body weight changes in NuTu-19 tail-vein injection rats. The results are shown as the mean ± SD for the tumor-forming group, non-tumor-forming group, and control group (* *P* < 0.05).

The pathological changes in the 4 tumor-forming rats and the 4 non-tumor-forming rats on day 60 are shown in Figure 3. On day 60, all tumor-forming rats showed severe lung metastatic nodules that scattered throughout the lung parenchyma. The lung tissue and the alveolar structure were showed severe destruction. The non-tumor-forming rats on day 60 did not show metastatic tumor nodes but had a large number of lymphocytes surrounding the bronchus. The periodic pathological changes of NuTu-19 tail-vein injection rats are shown in Figure 4. The lung metastatic tumor nodules became apparent 45 days after tail-vein injection of NuTu-19 cells, indicating that days 12 and 27 are in the early stages of lung tumor formation.

**Figure 3.**
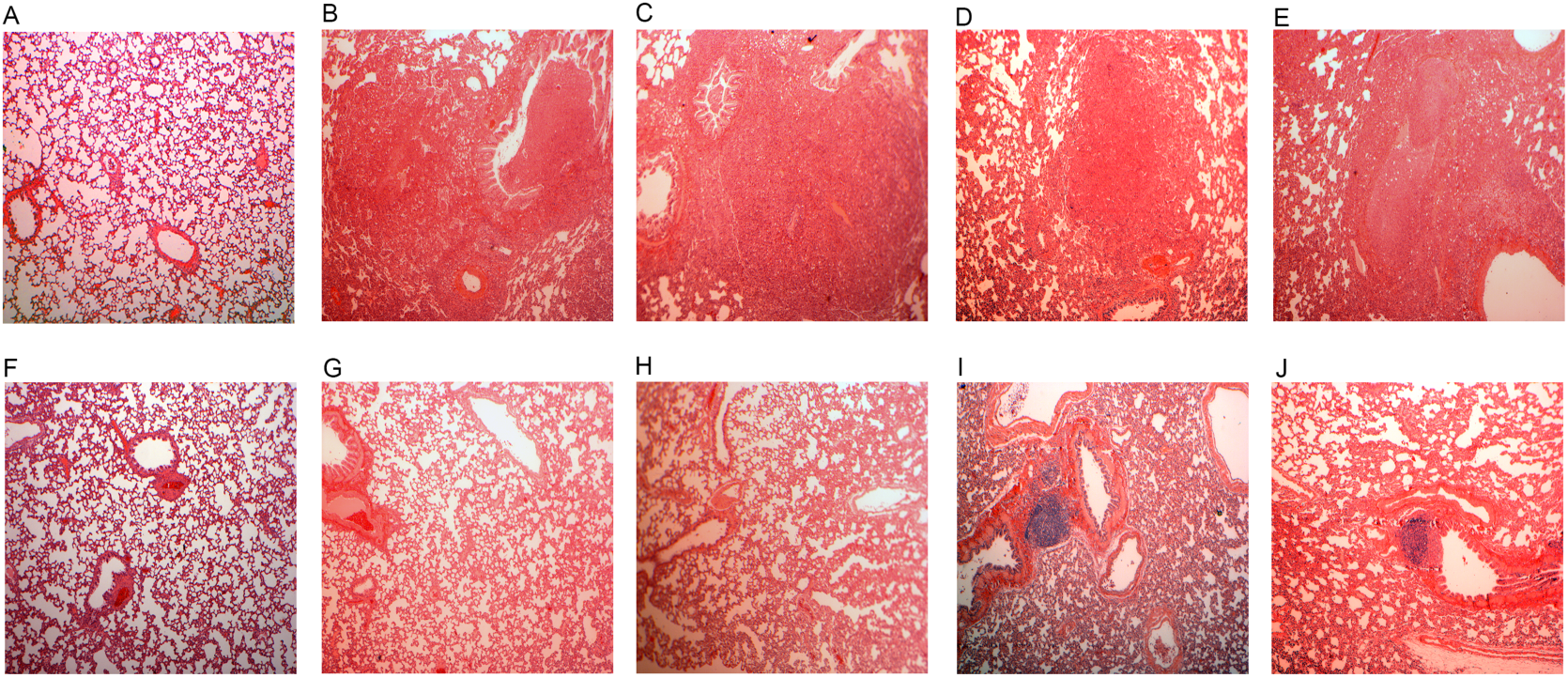
Pathological changes in NuTu-19 tumor-forming and non-tumor-forming rats on day 60. (A, F) The control group tail-vein injected with PBS. (B-E) Four NuTu-19 tumor-forming rats used for the discovery phase on day 60. (G-J) Four NuTu-19 non-tumor on day 60. The magnification was 40× for the images of H&E staining.

**Figure 4.**
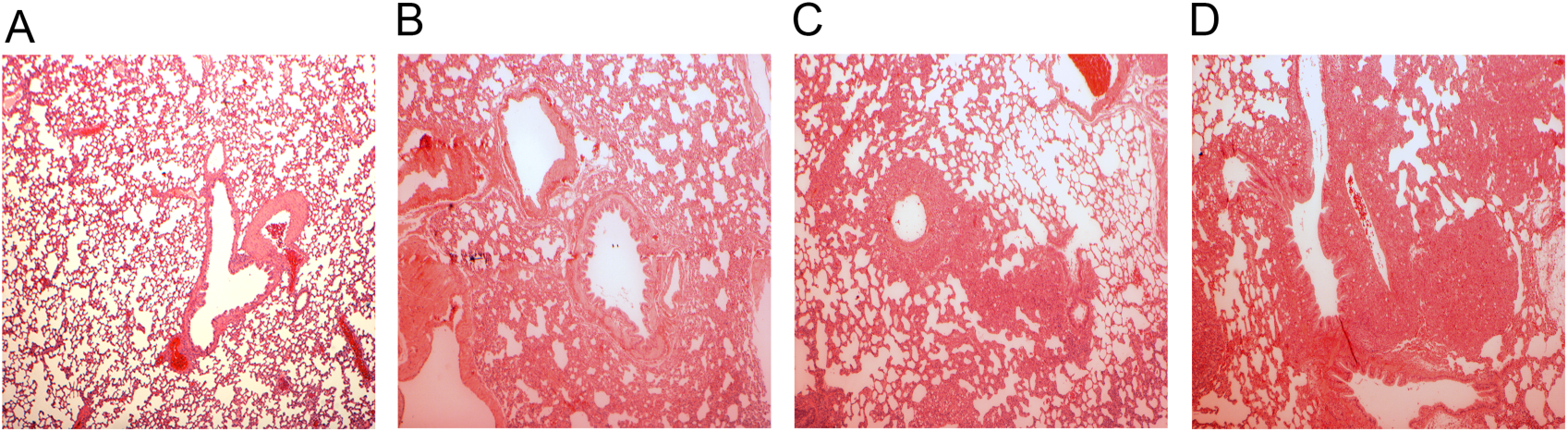
Pathological changes in NuTu-19 tail-vein injection rats. (A) The control group tail-vein injected with PBS. (B-D) NuTu-19 tail-vein injection rats on day 35, 45, 52. The magnification was 40× for the images of H&E staining.

### Urine proteome changes in tumor-forming and non-tumor-forming rats

Twenty urine samples from 4 tumor-forming and 4 non-tumor-forming rats at five time points (days 0, 12, 27, 39, and 52) were used for the discovery phase, with two technical replicates. Label-free LC-MS/MS quantitation was used to find the differential urinary proteins at these five time points. The quantification of tumor-forming rats was based on feature intensity using Progenesis software. A total of 532 proteins with at least 2 unique peptides were identified. The quantification of non-tumor-forming rats was based on spectra counts using the Scaffold software. A total of 505 proteins with at least 2 unique peptides were identified with < 1% FDR at the protein level. All identification and quantitation details are listed in Tables S3 and S4. By using screening criteria, 180 and 118 significantly changed proteins were identified in the tumor-forming and non-tumor-forming rats. Specifically, there were 99, 120,123,143 differential proteins on days 12, 27, 39, 52 in tumor-forming rats, respectively, while 34, 84, 91 and 96 differential proteins in non-tumor-forming rats. The details of these differential proteins at different time points are presented in Tables S5 and S6.

### Functional analysis

According to previous research, even when millions of cancer cells are injected into an experimental mouse, they might give rise to only a few metastases that are detectable in ‘end-point’ assays, indicating that metastasis is an inefficient process[20]. To determine why some rats formed lung tumors while others did not on day 60, functional annotation of differential proteins was performed using both DAVID and IPA. Specific biological processes, pathways and disease/functions were compared in tumor-forming and non-tumor-forming rats. Representative lists are presented in Figure 5. The cell component and molecular function enrichment analyses are shown in Figure S3 and S4. All these representative lists use a significance threshold of P < 0.05.

**Figure 5.**
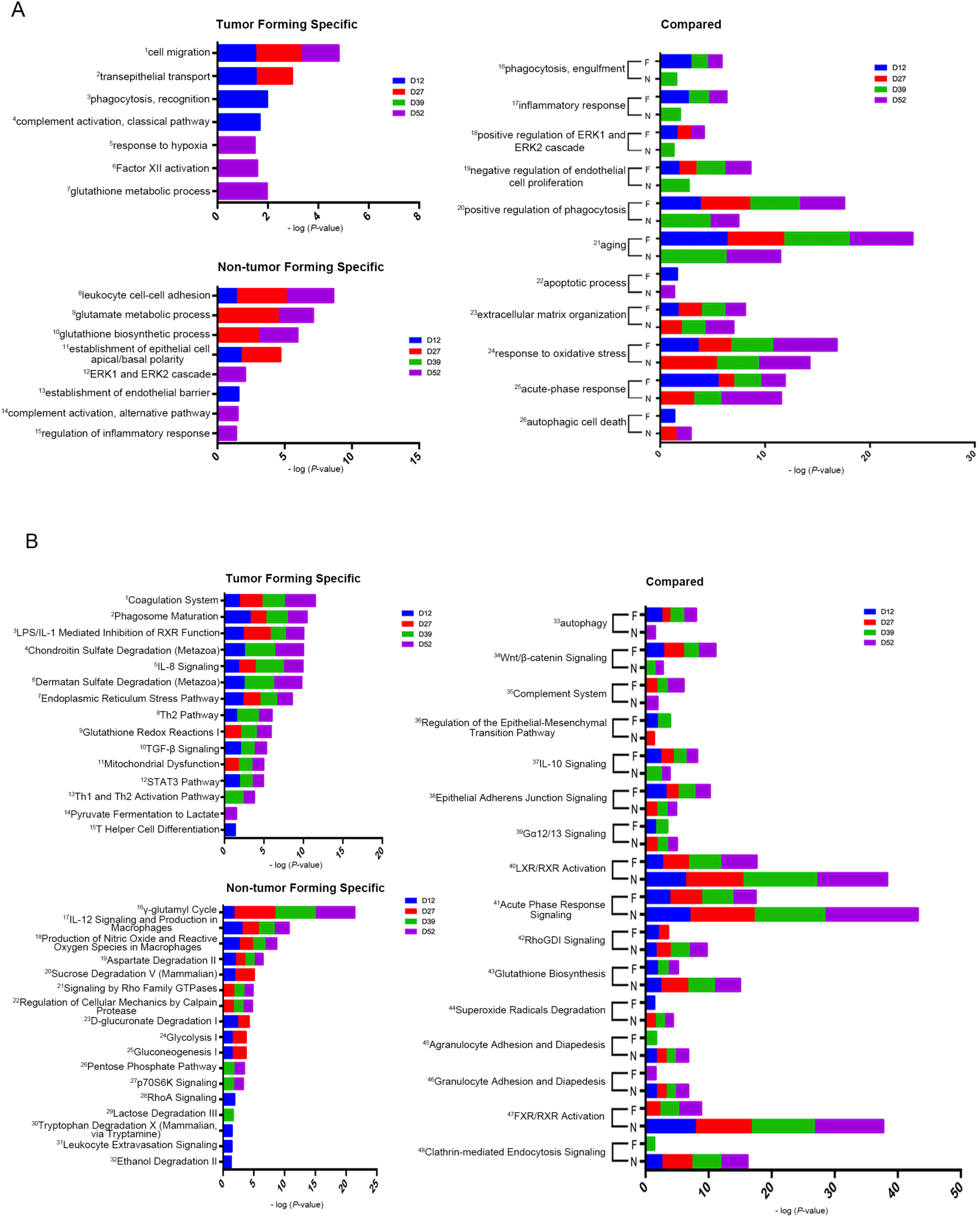
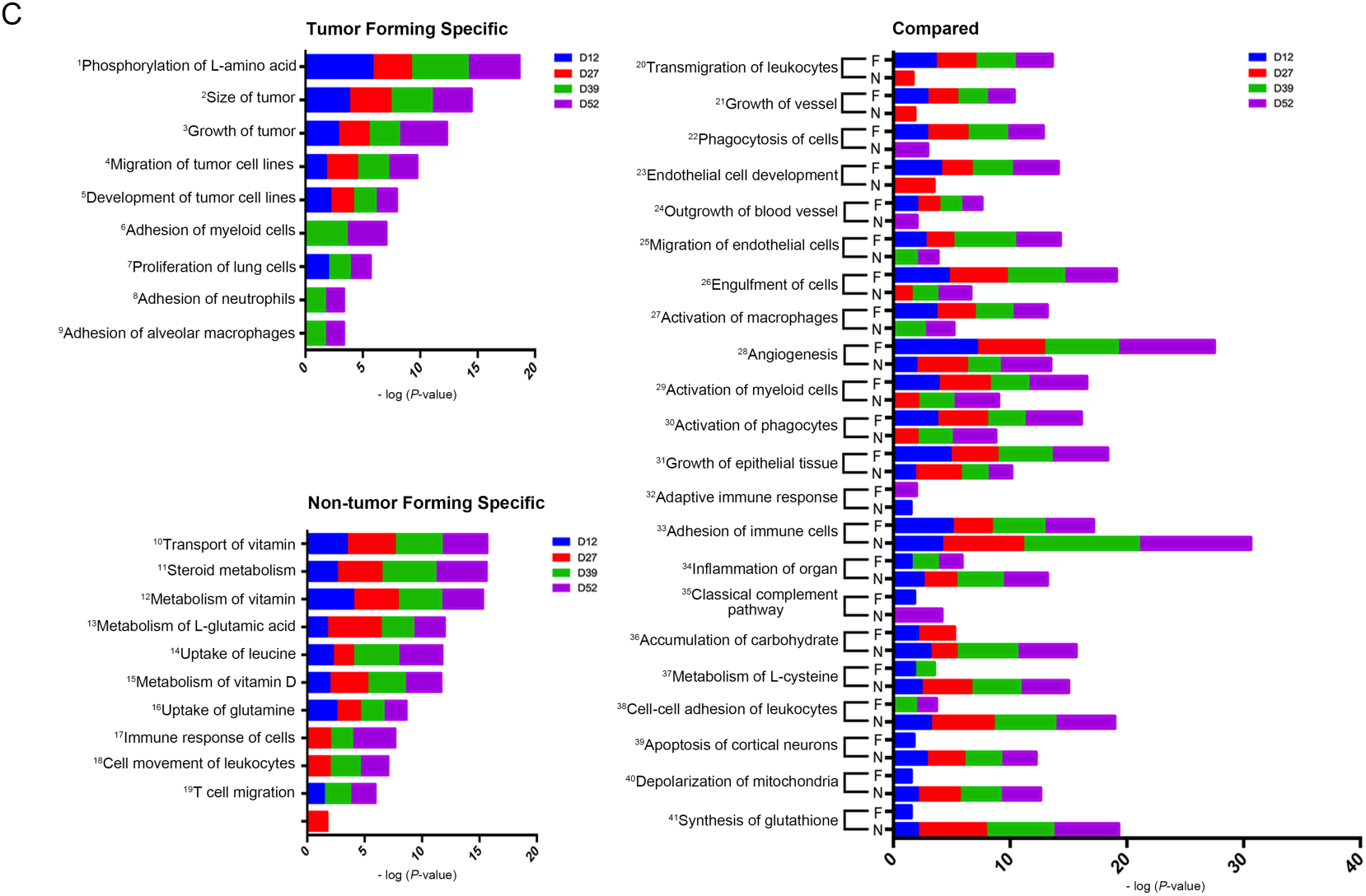
Functional enrichment analysis of tumor-forming and non-tumor-forming rats. Dynamic changes in the biological processes (A), pathways (B) and disease and functions (C) at multiple time points were classified. Each part was divided into tumor-specific, non-tumor-specific and compared analysis. All enrichment lists are with *P* < 0.05. F represents the tumor-forming group, and N represents the non-tumor-forming group.

Cancer metastasis involves a complex cascade of events: tumor cells i) invade the surrounding stroma and extracellular matrix; ii) intravasate into the bloodstream and survive in the blood circulation; iii) arrest at distant organ sites; and iv) extravasate and invade into the parenchyma of distant tissues[21, 22]. Some enriched functional lists were found to be associated with these cascades during this process and to be present only in tumor-forming rats. The enriched cell migration (Fig. 5A1) and migration of tumor cell lines (Fig. 5C1) are associated with the cancer cell intravasation process. When cancer cells enter the bloodstream, their interaction with other cell types can help them avoid immune attack and facilitate their passage to and extravasation at distant sites[23]. For example, i) the enriched coagulation system (Fig. 5B1) was reported to provide an additional layer of immune evasion, which contributes to tumor cell metastasis[21, 23, 24]; ii) TGF-β signaling (Fig. 5B10) was reported to be derived from platelets, and the adhered platelets can prevent tumor cell recognition and lysis by NK cells[25]; iii) the adhesion of neutrophils (Fig. 5C8) will entangle tumor cells in the circulation, which will make the tumor cells more apt to survive intraluminally, adhere to endothelial cells, and extravasate[26]; iv) enriched phagosome maturation (Fig. 5B2) and phagocytosis recognition (Fig. 5A3) have been reported to play a dichotomous role in cancer, where they promote tumor growth but also serve as critical immune effectors of therapeutic antibodies[27]; v) factor XII activation (Fig. 5A6) was reported to interact with monocyte/macrophages, which mediate peritoneal metastasis of epithelial ovarian cancer [28]. The enriched transepithelial transport (Fig. 5A2) and the adhesion of myeloid cells and alveolar macrophages (Fig. 5C6, C9) was reported to be associated with the extravasation process, which means that the carcinoma cells traverse the endothelial wall[29]. Other enriched functional lists were also associated with tumor progression. For example, i) the enriched functional of tumor size, tumor growth and tumor cell line development was associated with progression (Fig. 5C2, 3, 5); ii) the complement activation classical pathway (Fig. 5A4), IL-8 signaling, Th2 pathway, STAT3 pathway, Th1 and Th2 activation pathway and T helper cell differentiation (Fig. 5B5, 8, 12, 13, 15) were all accompanied by tumor progression[30–34]; iii) alterations in the phosphorylation of the L-amino acid STAT3 pathway (Fig. 5C1) were reported to result in serious outcomes in the form of diseases, especially cancer[35]; iv) the response to hypoxia STAT3 pathway (Fig. 5A5) induces the endoplasmic reticulum stress pathway, which was reported to be a major hallmark of the tumor microenvironment that is strictly associated with rapid cancer progression and the induction of metastasis[36–38]; v) the mitochondrial dysfunction STAT3 pathway (Fig. 5B11) was reported to play a critical role in cancer progression and metastasis[39, 40]; and vi) the pyruvate fermentation STAT3 pathway (Fig. 5B14) to lactate was reported as part of the Warburg effect, which benefits cancer cells[41, 42].

Some enriched functional lists were observed in both tumor-forming and non-tumor-forming rats but exhibited a stronger tendency in tumor-forming rats. Examples include i) the enriched phagocytosis engulfment, positive regulation of phagocytosis (Fig. 5A16, 20), phagocytosis of cells, engulfment of cells and the activation of phagocytes (Fig. 5C22, 26, 30), ii) the positive regulation of ERK1 and ERK2 cascade (Fig. 5A18), autophagy, Wnt/β-catenin Signaling, complement system, IL-10 signaling (Fig. 5B33, 34, 35, 37), activation of myeloid cells and activation of macrophages (Fig. 5C29, 30), and iii) the regulation of the epithelial-mesenchymal transition pathway, epithelial adherens junction signaling (Fig. 5B36, 38), growth of vessel, endothelial cell development, outgrowth of blood vessel, migration of endothelial cells, angiogenesis and the growth of epithelial tissue (Fig. 5C21, 23, 24, 31). All these lists were more associated with cancer metastatic progression.

Some enriched functional lists exhibited a strong association specifically in non-tumor-forming rats. For example, i) the glutamate metabolic process, glutathione biosynthetic process (Fig. 5A9, 10), γ-glutamyl cycle, aspartate degradation II (Fig. 5B16, 19), metabolism of L-glutamic acid, uptake of leucine and uptake of glutamine (Fig. 5C13, 14, 16) are associated with cancer glutamine metabolism. It has been reported that glutamine metabolism is essential for the proliferation of most cancer cells, and some cancer cells die if they are deprived of glutamine. The restriction of glutamine metabolism has been proven to be effective in inhibiting tumor growth both in vivo and in vitro through inducing apoptosis, growth arrest and/or autophagy[43]. Glutamine is first converted into glutamate via glutaminase (GLS/GLS2), while glutamate contributes to the synthesis of glutathione[44, 45]. Therefore, we believe that the items mentioned above will contribute to the non-tumor-forming process. ii) The enriched IL-12 signaling and production in macrophages (Fig. 5B17) were reported to activate all major cytotoxic killer and helper cell types of the immune apparatus (NK, NKT, CD4+, and CD8+ T cells), which are crucially important for immunosurveillance of and resistance to cancer development and progression. The extraordinary antitumor efficacy of IL-12 has also been demonstrated in animal models of cancer of diverse types, and its use in various forms is now involved in a large number of human cancer clinical trials[46]. iii) The production of nitric oxide and reactive oxygen species in macrophages (Fig. 5B18) was reported to be proinflammatory mediators that participate in the innate immune response[47, 48]. iv) The enriched signaling by Rho family GTPases and RhoA signaling (Fig. 5B21, 28) was reported to play an important role in pathological processes, including cancer progression, inflammation and wound repair[49]. v) The transport of vitamin D, steroid metabolism, metabolism of vitamin D, metabolism of vitamin D and steroid metabolism (Fig. 5C10, 11, 12, 15) were reported to be helpful in the treatment of cancer, especially when vitamin D metabolites or analogs were added to existing therapies[50].

Some of the commonly enriched lists were more prominent in non-tumor-forming rats. For example, i) enriched glutathione biosynthesis, superoxide radicals degradation (Fig. 5B43, 44), accumulation of carbohydrate, metabolism of L-cysteine, depolarization of mitochondria and the synthesis of glutathione (Fig. 5C36, 37, 40, 41), ii) agranulocyte adhesion and diapedesis (Fig. 5B45), granulocyte adhesion and diapedesis (Fig. 5B46), cell-cell adhesion of leukocytes (Fig. 5C38), iii) RhoGDI signaling, clathrin-mediated endocytosis signaling (Fig. 5B42, 48), iv) adhesion of immune cells, and organ inflammation (Fig. 5C33, 34).

Some additional enriched functional lists also deserved attention. For example, the enriched apoptotic process (Fig. 5A22) and the classical complement pathway (Fig. 5C35) were overrepresented in tumor-forming rats only on day 12 and overrepresented in non-tumor-forming rats only on day 52. In contrast, the enriched adaptive immune response (Fig. 5C32) was overrepresented in tumor-forming rats only on day 52 and overrepresented in non-tumor-forming rats only on day 12.

### Parallel reaction monitoring (PRM) validation

In the discovery phase, candidate protein biomarkers were identified using the label-free quantitative proteomics method. Sixty-two common differential proteins identified in tumor-forming and non-tumor-forming rats were excluded from the subsequent PRM analysis (Figure S5). All differential proteins identified on days 12 and 27 in tumor-forming and non-tumor-forming rats were validated by PRM (Table S7, S8).

For the tumor-forming group, 91 targeted differential proteins were used for PRM validation in another four tumor-forming rats. A total of 68 proteins were successfully quantified, while 38 of them were statistically significant. The following criteria were used for further selection: i) the same fold-change trend with label-free quantification compared to day 0, ii) identification on day 12 or day 27, and iii) the existence of a human orthologue. Finally, 22 urinary proteins changed significantly during the early phase of lung tumor formation. (Figure 6) Some of these differential proteins have already been reported to be associated with lung cancer biomarkers/pathology/mechanism or ovarian cancer metastasis. Several examples are provided below. (i) Insulin-like growth factor-binding protein 3 (IBP3) was reported as a lung cancer biomarker in serum, bronchoalveolar lavage fluid or lung tissue[51–53], and IBP3 was also reported to modulate lung tumorigenesis[54] and increase lung cancer risk in smokers[55]. In epithelial ovarian cancer, IGFBP-3 was reported to play an important role as an invasion-metastasis suppressor[56]. (2) Neuropilin-1 (NRP1) was reported to be highly expressed in NSCLCs, which play a key role in the occurrence, development and metastasis of NSCLC[57]. The increased levels of neuropilin-1 may be due to miR-152 suppression, suggesting a new therapeutic application of miR-152 in the treatment of NSCLC[58]. The coexpression of NRP1 and NRP2 genes was reported to be significantly correlated with tumor progression through neovascularization in NSCLC[59]. In addition, NRP1 was also reported as a valuable prognostic marker and a potential molecular therapy target for ovarian cancer patients[60]. (3) Tyrosine-protein kinase Mer (MERTK) was reported to be overexpressed in human non-small cell lung cancer[61]. (4) Sortilin (SORT) was reported as a regulator of EGFR intracellular trafficking, which in turn impacts lung tumor growth[62]. It was also reported that the silencing sortilin expression in tumor cells may introduce sortilin as a potential candidate for developing a novel targeted therapy in patients with ovarian carcinoma[63]. (5) Complement C4 (CO4) was reported as a lung cancer biomarker for diagnosis and prognosis[30, 64]. (6) Upregulation of Syndecan-4 (SDC4) was reported to contribute to the TGF-β1-induced epithelial to mesenchymal transition in lung adenocarcinoma A549 cells[65]. Other differentially expressed proteins are listed in Table 1.

**Table 1:**
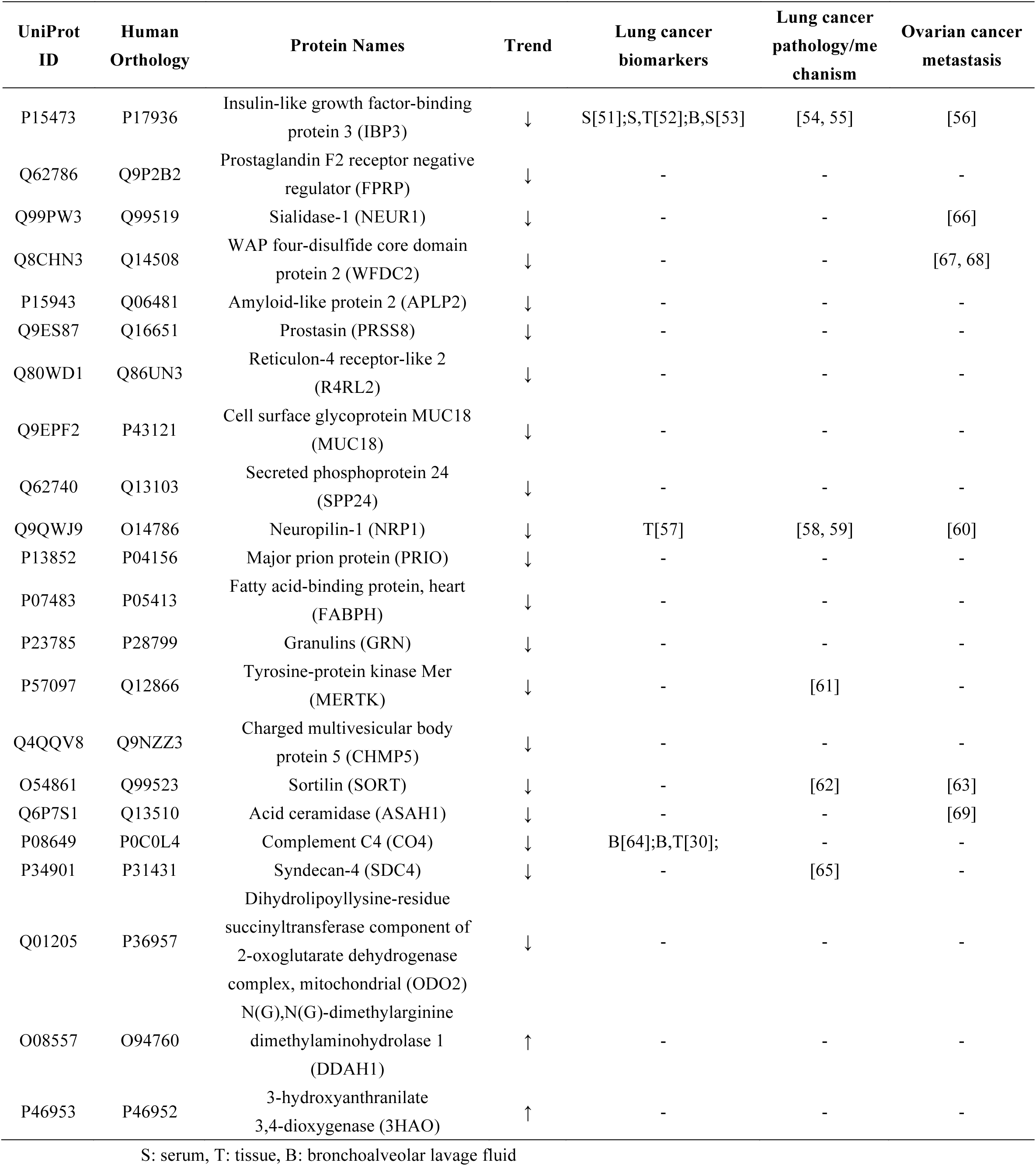
Candidate biomarkers for the early detection of lung tumor formation. After PRM validation in another four lung tumor-forming rats, the candidate differential proteins were searched in PubMed for lung cancer biomarkers/pathology/mechanism or ovarian cancer metastasis reports. S: serum, T: tissue, B: bronchoalveolar lavage fluid.

**Figure 6.**
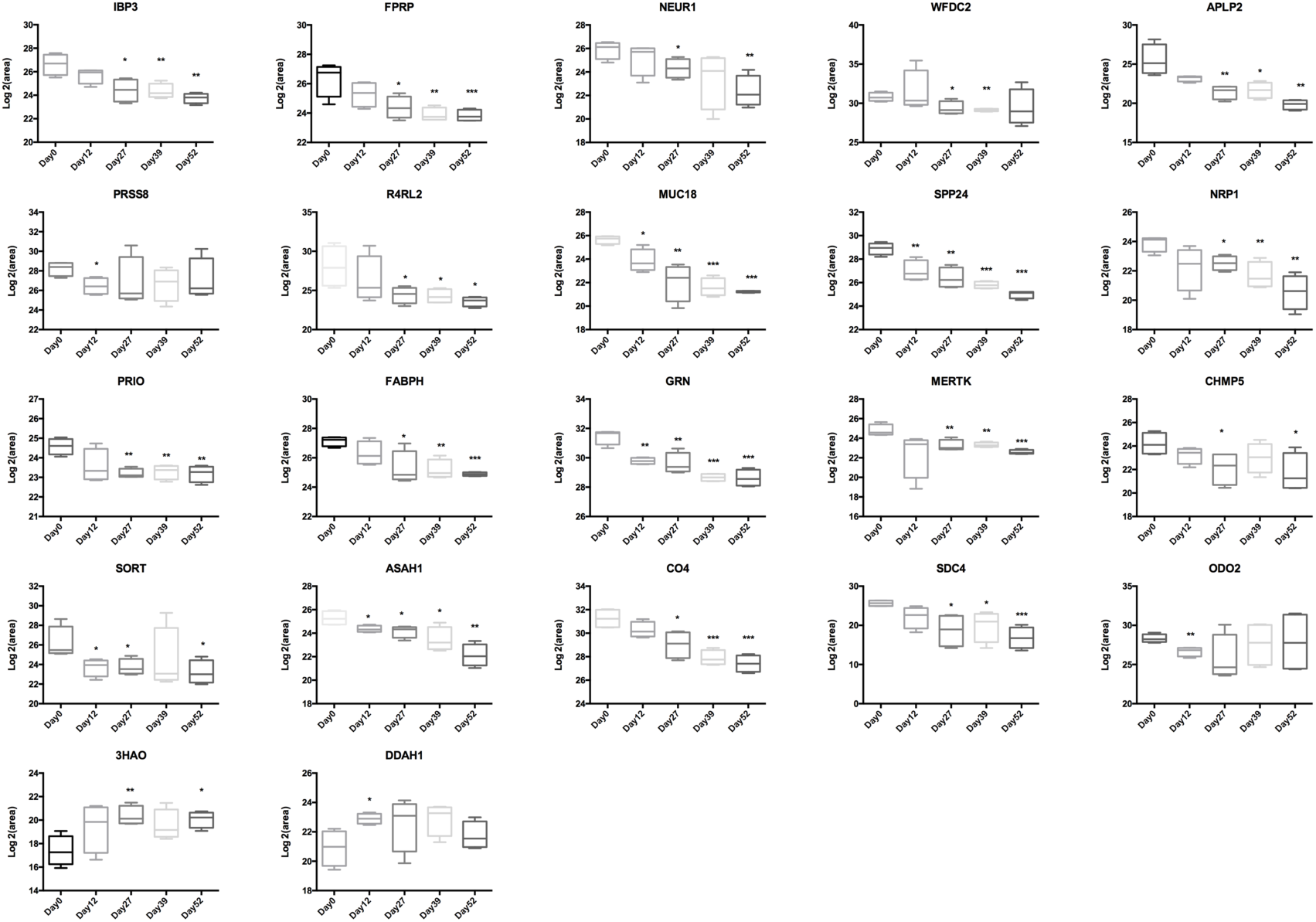
Expression of candidate urine biomarkers from NuTu-19 tumor-forming rats by PRM quantification. The x-axis represents different stages after the tail-vein injection of NuTu-19 tumor cells. The y-axis represents the log2 area of intensity based on PRM quantification. (*p<0.05, **p<0.01, ***p<0.001)

For the non-tumor-forming group, 44 targeted differential proteins were used for PRM validation in another three tumor-forming rats. A total of 41 proteins were successfully quantified, while 20 of them were statistically significant. After further selection with previous criteria, 14 of them were finally chosen for the early detection of non-tumor-forming processes in the lung. (Figure 7)

**Figure 7.**
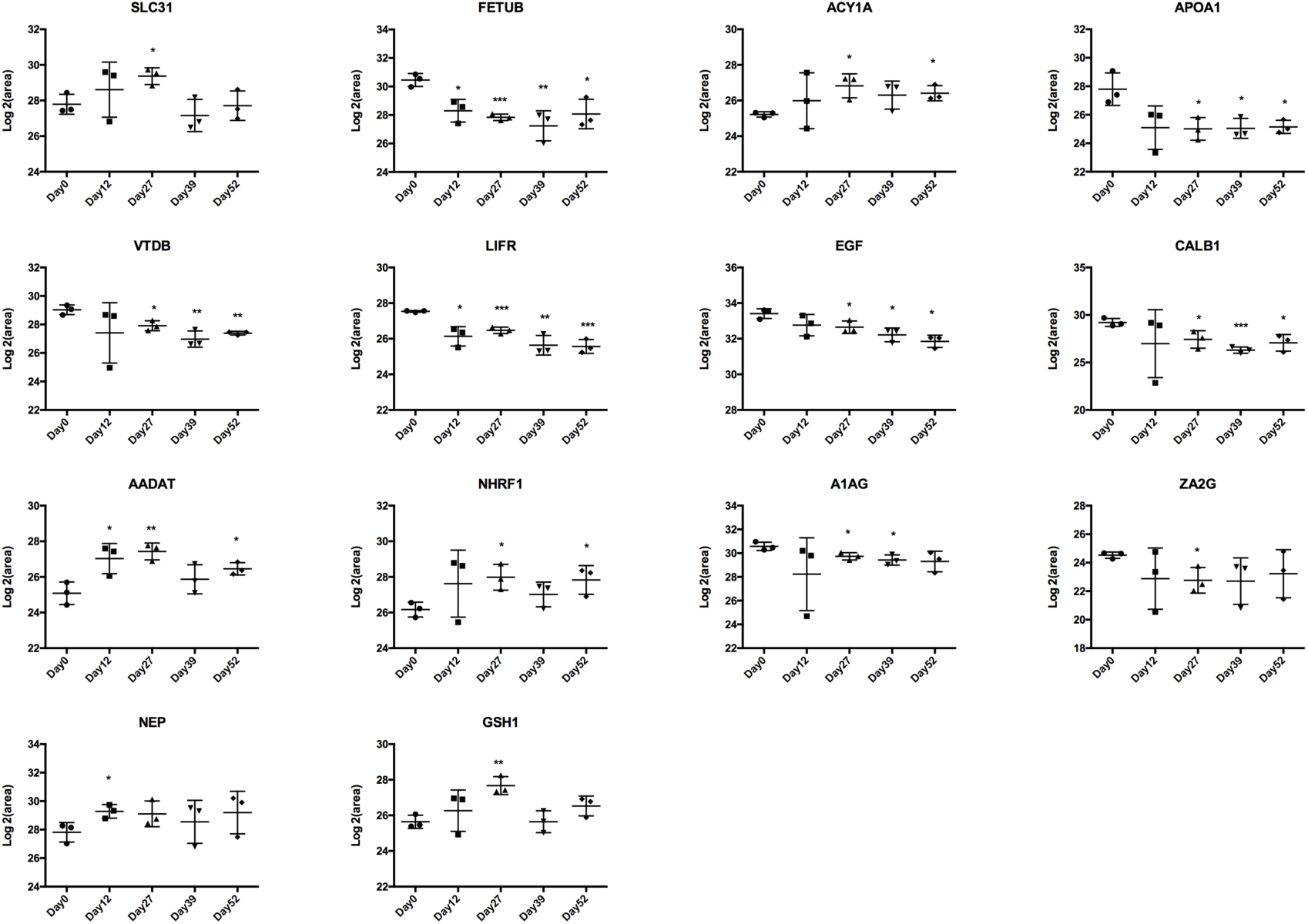
Expression of candidate urine biomarkers from NuTu-19 non-tumor-forming rats by PRM quantification. The x-axis represents different stages after the tail-vein injection of NuTu-19 tumor cells. The y-axis represents the log2 area of intensity based on PRM quantification. (*P<0.05, **P<0.01, ***P<0.001.)

## Conclusion

Overall, this work was a preliminary study in NuTu-19 tumor-forming and non-tumor-forming rats. Our results revealed that the urine proteome has the potential to differentiate tumor-forming and non-tumor-forming status in the lung at an early stage, which provides potential clues for early clinical intervention clinically. In future studies, the urinary protein biomarkers require further evaluation in a large number of clinical samples to test their sensitivity and specificity. Some significant pathways identified in non-tumor-forming rats in this study may also have potential applications in monitoring cancer treatment and prevention studies.

## Acknowledgments

National Key Research and Development Program of China (2018YFC0910202, 2016YFC1306300), Beijing Natural Science Foundation (7172076), Beijing Cooperative Construction Project (110651103), Beijing Normal University (11100704), Peking Union Medical College Hospital (2016-2.27).

## Supplementary Information

**Supplementary Table S1:** Retention times of 78 differential proteins with 237 peptides used for tumor-forming PRM validation.

**Supplementary Table S2:** Retention times of 42 differential proteins with 205 peptides used for non-tumor-forming PRM validation.

**Supplementary Table S3:** Identification and quantitation details of the urine proteome identified in the tumor-forming group.

**Supplementary Table S4:** Identification and quantitation details of the urine proteome identified in the non-tumor-forming group.

**Supplementary Table S5:** Differential proteins identified on days 12, 27, 39, and 52 in the tumor-forming group.

**Supplementary Table S6:** Differential proteins identified on days 12, 27, 39, and 52 in the non-tumor-forming group.

**Supplementary Table S7:** Differential proteins identified on day 12 and day 27 specifically in the tumor-forming group.

**Supplementary Table S8:** Differential proteins identified on day 12 and day 27 specifically in the non-tumor-forming group.

**Supplementary Figure S1:** The CV values of 237 PRM-targeted peptides for tumor-forming validation.

**Supplementary Figure S2:** The CV values of 205 PRM-targeted peptides for non-tumor-forming validation.

**Supplementary Figure S3:** The cell component enrichment analysis of differential proteins identified in the tumor-forming group (A) and the non-tumor-forming group (B).

**Supplementary Figure S4:** The molecular function enrichment analysis of differentially expressed proteins identified in the tumor-forming group (A) and the non-tumor-forming group (B).

**Supplementary Figure S5:** Comparison of differential urinary proteins between the tumor-forming group and the non-tumor-forming group.

